# On the Regulator of Spike Activity in Cortical Neurons Under Hypothermic Conditions

**DOI:** 10.1101/041004

**Authors:** Yu. S. Mednikova, N. M. Zakharova, N. V. Pasikova, I. V. Averina

## Abstract

In sensorimotor cortical slices of guinea pig in the course of cooling incubating fluid from 34 to 21-22°C it was shown that hypothermia exerted both increase and decrease of spontaneous activity in different neurons. On hypothermic increase of firing level spike responses of soma to iontophoretic application of glutamate to dendritic locus appeared with shorter latencies and with longer latencies – on hypothermic decrease of spontaneous activity. At the same time hypothermia did not influence on the evoked spike reactions to iontophoretic application of glutamate straight to the soma. It means that hypothermic disorders of neuronal activity are not connected with changes in sensitivity to glutamate but determined by changes of amplitude of glutamatergic excitation while propagating along dendritic branches. The changes in spontaneous activity began at 30°C along with the decreased spike reactions to iontophoretic applications of acetylcholine and efficacy of dendro-somatic propagation. At the same temperature the fall of spike amplitude was initiated and increased with further hypothermia. It is proposed that the basis for hypothermic changes of neuronal activity is the decreased rate of M-cholinergic process at 27-29°C which leads both to attenuation of conductive function of dendrites and imbalance of K^+^ ion homeostasis. Peculiarities of hypothermic regulation of neuronal spike activity depend on individual functional properties of cortical neurons.

## 1. Introduction

When applying hypothermia and hypoxia exert almost identical disorders in nerve tissues and that is why hypothermic and hypoxic changes in neuronal activity are often considered together (Alkan, Korfali, 2000; Boutilier, 2001; Hochachka, 1986). During long periods of even slight and moderate hypothermia (*t*=32-34 and 28°C), as well as during hypoxic periods of any degree, in nerve cells membrane permeability to ions, supporting the stability of membrane potential, increases, resulting in a flow of Na^+^ and K^+^ in the direction of their thermodynamic gradients. The occurring decrease of membrane potential opens potential-dependent Ca^2+^ channels and leads to pathologic entry of Ca^2+^ into cytosol, activation of phospholipases and proteases, disintegration of neuronal membranes and necrotic death of the neurons. (Boutilier, 2001; Hochachka, 1986). Such a dramatic chain of events describes the hypoxia-related disorders quite well, because the first sign of hypoxic condition of the brain is a blockade of Na^+^-K^+^-ATPase activity caused by deficiency of energy supply (Lipton, 1999). But under hypothermic condition the reason for imbalance of ionic processes remains unclear: the activity of Na^+^-K^+^-ATPase even increases with fall of temperature (Aslanidi et al.,1997); membrane potential and spike amplitude in neurons of frogs living at low temperatures mortal for the brain of homoeothermic animals have the same values as in mammalian neurons (Ecceles, 1957), while the energetic demands of cold-blooded brain significantly less (Hochachka, 1986; Ivanov, 2004). That is why neither temperature-dependence nor energetic limitations of Na^+^-K^+^-ATPase activity should be considered as the reasons for the imbalance of ionic homeostasis in the cases of hypothermia. Noting this contradiction, P.W.Hochachka had supposed that there should be an additional mechanism sensitive to hypothermia, coupling membrane functions and cellular metabolism, which had not been taken into account earlier (Hochachka, 1986). The evidence for such a mechanism follows from the fact that a lot of hypothermic alterations of brain activity are linked to the same temperature point: below 28°C the mammals lose consciousness (Ivanov, 1996; Prosser, 1973), below 28° inflation of cortical EEG occurs (Ignat’ev et al., 2005), below 27-29°C the frequency of spontaneous neuronal activity decreases together with the temperature drop of the rate of M-cholinergic reaction (Mednikova et al.,2002; Mednikova et al.,2008).

M-cholinergic process is one of the main reactions of the brain. Its designation consists in regulating membrane properties of neurons via metabolic process limiting K^+^ permeability of cellular membranes, thus increasing their specific resistance (Krnjević, 1971; McCormick, Prince, 1986; Brown et al., 1995). High sensitivity of M-cholinergic reaction to temperature (Brown et al., 1995; Mednikova et al., 2008) and its dependence from energy supply (Godfraind et al., 1971) were established. Functional role of cholinergic process is related to regulation of conductive function of dendrites, providing the decrease of excitation decrement when excitation is propagating from dendrites to soma (Mednikova et al.,1998) and thus providing the increase of the level of spontaneous neuronal activity (Mednikova et al.,2010).

Taking into account that one of the parameters of neuronal activity regulated by hypothermia is spontaneous activity (Mednikova et al., 2002), we analyzed the variability of this parameter under hypothermic conditions in several groups of neurons depending on the initial activity level, dendrosomatic conductance factor and the expression of temperature-dependent M-cholinergic reaction. We hope the studying parameters lead us to mechanisms of hypothermic disorders of brain activity.

## 2. Methods

### 2.1. Slice preparation and incubation conditions

Animals were treated with observance of recommendation on ethics of work with animals offered by European Communities Council Direction (86/609 EEC) and experimental protocols approved by ethics committees of Institute of Higher Nervous Activity Russian Academy of Sciences. The experiments were carried out on slices of sensorimotor cortex of guinea pigs (weight 200-250 g). After fast decapitation of the animals by guillotine and scull break-up, the brain surface was sprinkled by ice-cold aerated Ringer-Krebs solution. 500 μm thick slices were prepared from longitudinal cortex block on VSL vibratome (World Precision Instruments, USA). Incubation chamber, where the slices were placed, consisted of two cells (reserve and experimental) with independent flow of Ringer-Krebs solution. Incubation solution saturated by gas mixture (95%O_2_ +5%CO_2_) consisted of the following components (mM): 124 – NaCl; 5 – KCl; 1.24 – KH2PO4; 1.3 – MgSO_4_; 2.4 – CaCl_2_; 26 – NaHCO_3_ and 10 – glucose (pH 7.4). Flow rate of incubating solution was 1.5-3 ml/min. The slices were incubated at room temperature for half an hour after preparation. Then, the temperature was increased *32-34*°C. The slices were incubated at this temperature for 1.5-2 hours before the beginning of spike activity registration. The pre-heating of incubation medium was performed by U1 thermostat (VEB, Germany). For finer temperature regulation, a Peltier element-based thermostating device was used (NPO «Biopribor», Russia). The temperature in experimental cell was controlled continuously by an electronic thermometer (NTC “NIKAS”, Russia)

### 2.2. Spike activity registration and transmitter microapplication.

#### 2.2.1. Electrodes and recording equipment

Three-channel glass microelectrodes (diameter 7.4-8 μm) were used for extracellular registration of neuronal spike activity and for iontophoretic transmitter application. The channel for registration and channel for controlling iontophoretic current were filled with 3M NaCl. The third channel contained 1M sodium glutamate (pH 7.5; Sigma Chemical Co., USA) or 2M acetylcholine chloride (pH 4.0; Sigma Chemical Co., USA). The channel for controlling iontophoretic current was often replaced by a second channel for transmitter phoresis, because the action of currents chosen for transmitter phoresis but being passed through current controlling channel did not evoke spike responses of recorded neurons. After amplification (DAM 80, World Precision Instruments, USA) and digitalization (E14-440, L-Card, Russia) the spike activity was inputted into computer (Intel (R) Core (TM) Duo) for storage, reproduction and processing of registered signals.

#### 2.2.2. Registration area and iontophoretic current parameters

A three-barrel microelectrode containing channel for recording was moved along V layer of the cortex (1.2-1.6 mm from pial surface) in order to find spike activity of neurons and transmitter applications to the soma of registered nerve cells. After the detection of spike activity of a neuron, the independent micropipette, containing only glutamate phoresis channel, was placed at different points of presumable dendritic tree. Short-time bursts of spike activity from “dendritic electrode” during glutamate application served as indicators of its location near dendritic surface (Mednikova, Karnup, 1995). The distance between somatic and dendritic micropipettes was defined with an eyepiece provided with micrometric scale. Glutamate was applied to dendrites and soma by 80 nA current (negative pole inside the electrode); acetylcholine was applied to soma by 70 nA current (positive pole inside the electrode). The duration of iontophoretic current for glutamate application was from 1 to 4.5 s, whereas for acetylcholine application it was always 4.5 s. Throughout an intervals between iontophoretic ejections the retaining 3-5 nA current of opposite direction was set in each of phoresis channel. Iontophoretic glutamate application to local dendritic points in the course of gradual withdrawal of micropipette from the point of maximal effect allowed to find out that the effective dose of glutamate near dendritic surface could be achieved at the distance not exceeding 20 μm, which implies a high locality of iontophoretic action.

### 2.3. Experimental protocols

Standard temperature, 32-34°C, was maintained constantly in the reserve cell of incubating chamber, whereas in the experimental cell it was maintained during the procedure of searching for spike activity of the neurons and its control testing. Cooling of the solution in the experimental cell to 21-24°C was performed with the rate of two degrees per minute. The rate of temperature restoration was the same. Thus, the whole cycle of hypothermic action took not longer than 10 minutes. During cooling, the neurons were always tested by a certain application manner: glutamate application to soma or dendrites every 12 s or acetylcholine application to soma every 24 s. If registration conditions allowed, the neuronal activity was observed during an hour after the episode of hypothermia. Every hypothermic action was carried out usually on a fresh slice in order to avoid post-hypothermic events.

### 2.4. Data analysis

The parameters of spike activity were analyzed by Power-Graph 3.3 computer program (PO «Power-Graph», Russia). An average level of spontaneous activity in 3-second interval before every iontophoretic application was calculated for pre-hypothermic, hypothermic and post-hypothermic periods. Spike amplitudes were also determined in these periods. Spike reactions of the neurons to transmitter application were estimated by duration of latency period of the response and its intensity. Difference between maximal current average of spike frequency in the response and in preceding background activity served as estimation of the intensity of the reaction. The significance of the parameters’ alterations was determined by non-parametric statistical methods (Sachs, 1972).

## 3. Results

### 3.1. Effect of hypothermia on spike frequency and spike amplitude

In the experiments on the layer V of sensorimotor cortex slices, the activity of 111 neurons was registered. The level of spontaneous activity varied from 0 to 25 impulses per second in different neurons. The distribution of the neurons by frequency of spontaneous activity is presented in the table.

**Table.** Distribution of 111 neurons of sensorimotor cortex according the level of spontaneous activity

| Level of spontaneous activity (imp/s) | 0 | 0–4 | 4–8 | 8–12 | 12–16 | 16–20 | 20 and more |
| --- | --- | --- | --- | --- | --- | --- | --- |
| Number of neurons | 42(38%) | 27(24%) | 21(19%) | 9(8%) | 4(4%) | 5(4%) | 3(3%) |

It is seen that a significant majority of the neurons is lacking the background activity. Presence of such neurons was detected by arising of spike activity in response to glutamate applied to the soma by short pulses of current. Among 111 nerve cells, 37.8% (42 neurons) had no spontaneous activity, 27 neurons (24.4%) had spontaneous activity frequency up to 4 impulses per second, whereas high-frequency neurons (8-25 impulses per second) comprised 18.9 % of the population (see the table). The results of previous experiments (Mednikova et al.,2008; Mednikova et al., 2010) and data obtained by other authors (Pennartz et al., 1998) gave evidence that alteration degree of spontaneous activity of the neurons significantly depends on their initial firing level. That’s why, the analysis of spontaneous activity frequency under short-time hypothermia was performed separately for the neurons divided in groups depending on the background activity level. Group I included neurons without background activity; group II, – the neurons with activity below 4 impulses per second; group III, – neurons with activity from 4 to 8 impulses per second; group IV, – high-frequency neurons (from 8 to 25 impulses per second)

Overall, 35 neurons were registered during hypothermia procedure. Figure 1 shows most typical cases of change of spontaneous activity level in the neurons of all four groups during decrease of incubation liquid temperature in experimental cell from 32-34 to 21-22°C (Fig.1, *A*) and further restoration of the temperature to the initial values (Fig.1, *B*).

**Fig. 1.**
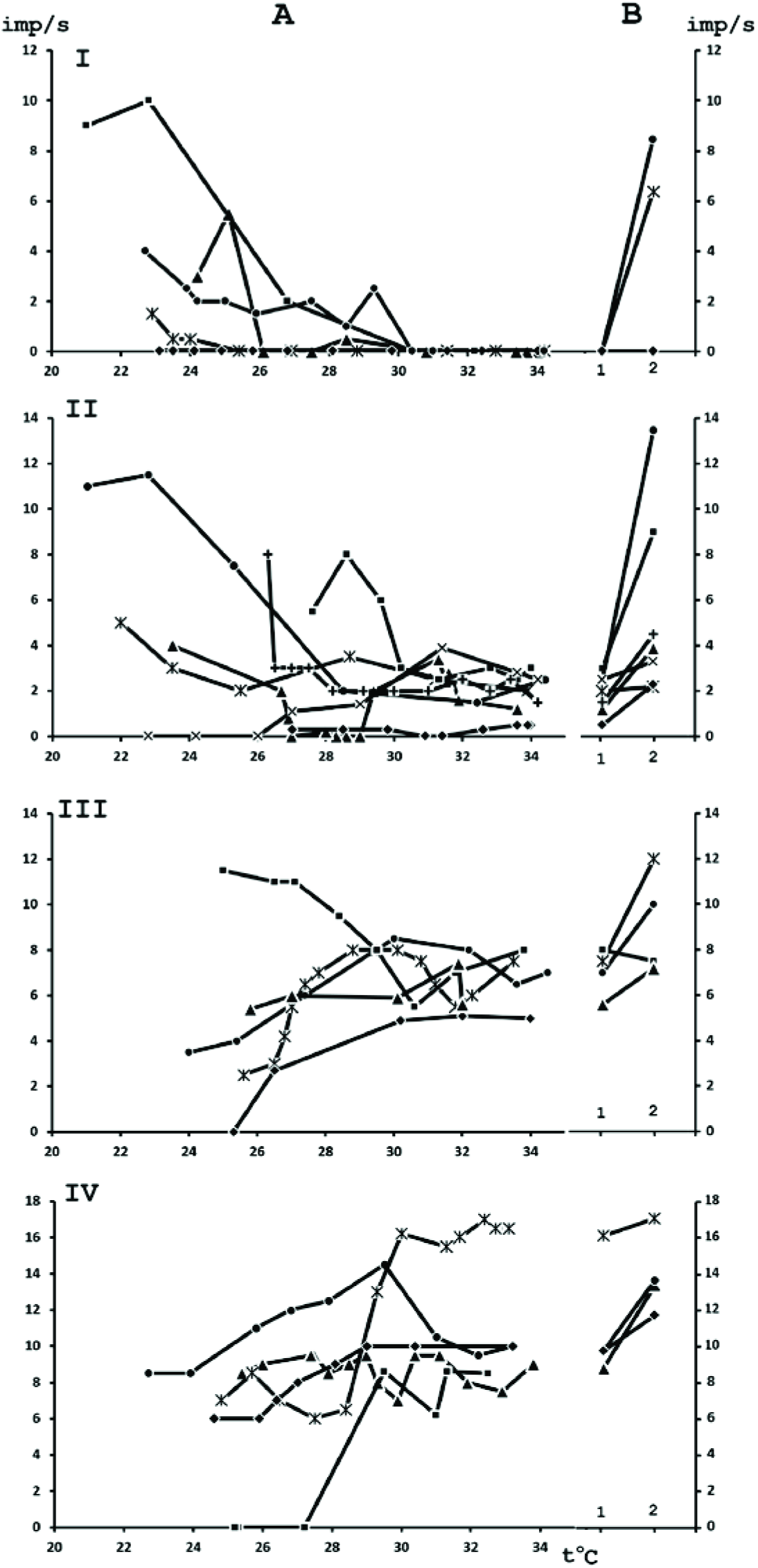
Frequency of spontaneous activity of sensorimotor cortex neurons in the course of decreasing temperature from 34 to 21°C and after restoration of the initial temperature values. Each graph illustrates the dynamics of single neuron spontaneous activity frequency during cooling from 32-34° to 21-22°C (A) and after restoration of the initial temperature (B). I, II, III, IV – four groups of nerve cells with different levels of spontaneous activity before cooling: I – neurons without spontaneous activity (0 imp/s); II – neurons with spontaneous activity frequency up to 4 imp/s; III – neurons with spontaneous activity frequency from 4 to 8 imp/s; IV – neurons with spontaneous activity frequency above 8 imp/s. A: abscissa – temperature, ordinate – spontaneous activity frequency, (imp/s). B: horizontal axis - 1 and 2, - the same temperature value before (1) and after (2) applying hypothermia; vertical axis – spontaneous activity frequency, (imp/s).

The specific feature for neurons of all four groups was the lack of significant difference in spontaneous activity level at 30°C compared to 34°C (Wilcoxon matched pairs signed rank test, α>>5%). Systematic changes of neuronal activity frequency began to appear by cooling incubation medium below 30°C, having different rate of firing change in different neurons and as it quite interesting, these changes were both increase and decrease of spike activity in different neurons. The neurons with the same level of spontaneous activity under cooling and at *t*=32-34°C were also present. Despite all the diversity of happening alterations, their dependence from the initial spontaneous activity level before cooling can be seen. Spontaneously inactive neurons almost in all cases of temperature fall below 30°C displayed the increase of spontaneous activity frequency the more significant the deeper cooling (Fig.1, *A,I*). Among the neurons with spontaneous activity up to 4 impulses per second, the increase of activity rate during cooling also prevailed, but the decrease of spontaneous activity could also be found in certain neurons (Fig.1, *A,II*). In the group of neurons having the activity of 4-8 impulses per second, the majority of registered cells displayed the decrease of spontaneous activity below 30°C, but one neuron showed growth of frequency (Fig.1, *A,III*). Finally, high-frequency neurons showed only decrease of spontaneous activity under hypothermia (Fig.1, *A, IV*). The neurons from groups III and IV showed significant decrease of spontaneous activity under cooling revealed at t=26°C compared to the level at 30°C (Wilcoxon matched pairs signed rank test, α<1%). The significant increase of activity rate under hypothermia in groups I and II was achieved at t=24°C compared to 30°C (Wilcoxon matched pairs signed rank test, α<1%).

Restoration of initial temperature values (32-34°C) after episodes of hypothermia lead to increase of spike activity compared to the level before cooling in almost all neurons independent of the character of activity changes under cooling (Fig.1, *B*; Wilcoxon matched pairs signed rank test, α<0.1%). Hyperactivity of low-frequency neurons (groups I and II) was characterized by an increase of frequency by average 231.4±40% compared to the level before cooling, and for the neurons with activity above 4 impulses per second (groups III and IV) it was only by 34.9 ± 5% of the initial level. The hypothermia-induced increase of spike activity was maintained during long time after restoration of initial temperature: in one third of the neurons recovery period lasted as long as the post-hypothermic time of observation (1 hour). Fast restoration of activity frequency was detected mainly just for high-frequency neurons from group IV, which displayed full restoration of activity in 10-20 min after episodes of hypothermia.

Cooling-related alterations of spontaneous activity level in nerve cells were accompanied by decrease of spike amplitude observed in all neurons independently of the direction of change of neuronal activity level under hypothermia (Fig.2, *A*).

**Fig. 2.**
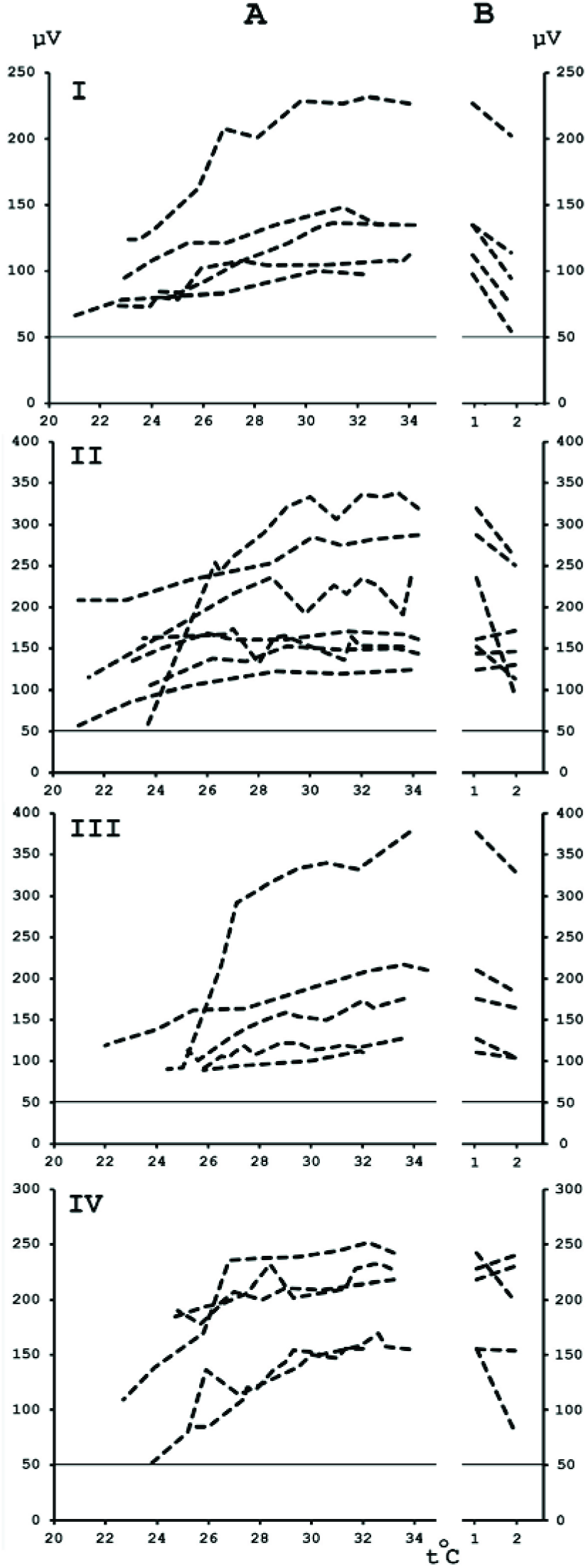
Spike amplitude in sensorimotor cortex neurons during decrease of temperature from 34 to 21 °C and further restoration of initial temperature values. Each graph illustrates the dynamics of single neuron spike amplitude changes during cooling from 32-34° to 21-22°C (A) and during restoration of the initial temperature (B). Temperature-related changes of spike amplitude presented for the neurons which spontaneous activity is indicated on Fig.1. I, II, III, IV – four groups of nerve cells with different levels of spontaneous activity before cooling (as shown on Fig.1). A: abscissa - temperature, ordinate – spike amplitude, μV. B: horizontal axis - 1 and 2, the same temperature value before (1) and after (2) applying hypothermia; vertical axis – spike amplitude, μV. The line parallel to abscissa axis on each fragment designates noise level of the recording equipment.

The decrease of spike amplitude during cooling of incubation medium from 34 to 30°C was very slight, equaling 6.3±0.5 μV in average (Wilcoxon matched pairs signed rank test, α<5%). A dramatic drop of spike amplitude, by 34.5±4 μV on average, occurred when cooling from 30 to 26°C (Wilcoxon matched pairs signed rank test, α<0.1%). Further cooling led to even more significant decrease of spike amplitude down to complete spike elimination in the noise of certain neurons (Fig.2, *A*). Restoration of initial temperature in overwhelming majority of the cases did not lead to total restoration of spike amplitude (Fig.2, *B*). The decrease of spike amplitude compared to the value before cooling was 15.5% on average (Wilcoxon matched pairs signed rank test,α<0.1%). Half of the neurons did not display full restoration of spike amplitude even in hour of observation after hypothermic episode. Only in the group of high-frequency neurons (group IV) the majority of neural cells showed restoration of initial spike amplitude right after restoration of initial temperature of incubation medium (Fig.2, *B, IV*).

### 3.2. Hypothermic attenuation of cholinergic reaction and regulation of spontaneous activity

Thus, the level of spontaneous activity and spike amplitude of the neurons undergo the most significant alterations in temperature range 27-29 °C. As shown earlier, a dramatic decrease of activation response to acetylcholine applied iontophoretically to nerve cells takes place in the same temperature range (Mednikova et al.,2008). The reaction to acetylcholine comprises slow increase in spike activity frequency during quite a long time (10 s and more) after termination of phoretic current (Fig.3, *1*), which shows that its development results from muscarinic mechanism (Krnjević et al.,1971; McCormick D.A., Prince D.A. 1986). Nine neurons tested by acetylcholine applications during hypothermic action showed no significant difference in spike rate increments over background activity in response to acetylcholine at 34 and 30 °C (Wilcoxon matched pairs signed rank test, α>>5%). Meanwhile, at t=26-27°C acetylcholine applications exerted either no increase of spike activity above background level or the increase of firing was much weaker and had a longer latency period (Fig.3, *3*). The reaction to acetylcholine estimated by increment of spike rate over the background level, was significantly lower for all 9 neurons at t=26°C than at t=30°C (Wilcoxon matched pairs signed rank test, α<1%). It is also clearly shown on Fig.3 that spontaneous activity of the neuron vanishes simultaneously with the weakening of the reaction to acetylcholine at 27°C (Fig.3, 2,3), which indicates on common temperature dependence of cholinergic process and the process of regulating spontaneous activity.

**Fig. 3.**
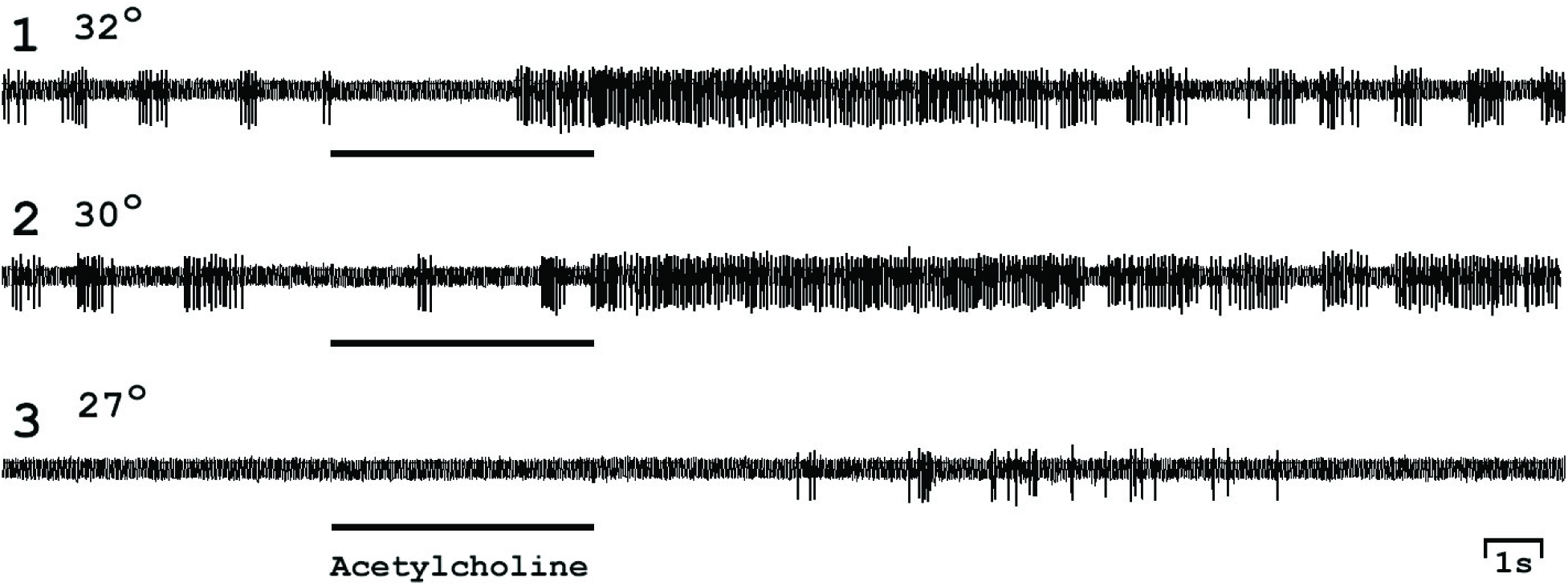
Responses of the sensorimotor cortex neuron to iontophoretic acetylcholine application during cooling of incubation medium. Acetylcholine was applied to the soma by 70 nA current (positive pole inside the electrode) simultaneously with recording spike activity of the neuron. Neuronal spike activity was recorded at following temperatures: 32°C(1), 30°C(2) and 27°C(3). Duration of acetylcholine electrophoresis current is shown as a line under each record of neuronal activity.

Fig. 4 graphically describes the changes of spontaneous activity level and frequency increment value under acetylcholine action during the decrease of temperature from 30-31 to 26-27°C. The temperature-related differences are presented for 9 neurons in percentage to the level of spontaneous activity (Fig.4,A) and the expression of response to acetylcholine (Fig.4,B) at *t=*30-31°C. As well as shown on Fig. 1, the level of spontaneous activity in different neurons was either decreasing (5 neurons), or increasing (4 neurons) at 26-27°C (Fig.4, A). At the same time, the response to acetylcholine during cooling only decreased independent from the character of spontaneous activity changes; for 7 of 9 neurons, the inhibition of the reaction was more than 60% (Fig.4, *B*). Thus, the decrease of M-cholinergic reaction degree in temperature range 27-29°C causes both increase and decrease of spontaneous activity frequency in different neurons. This means that the result of temperature-dependent decrease of cholinergic reaction rate is action on two different mechanisms regulating spontaneous activity level in opposite directions.

**Fig. 4.**
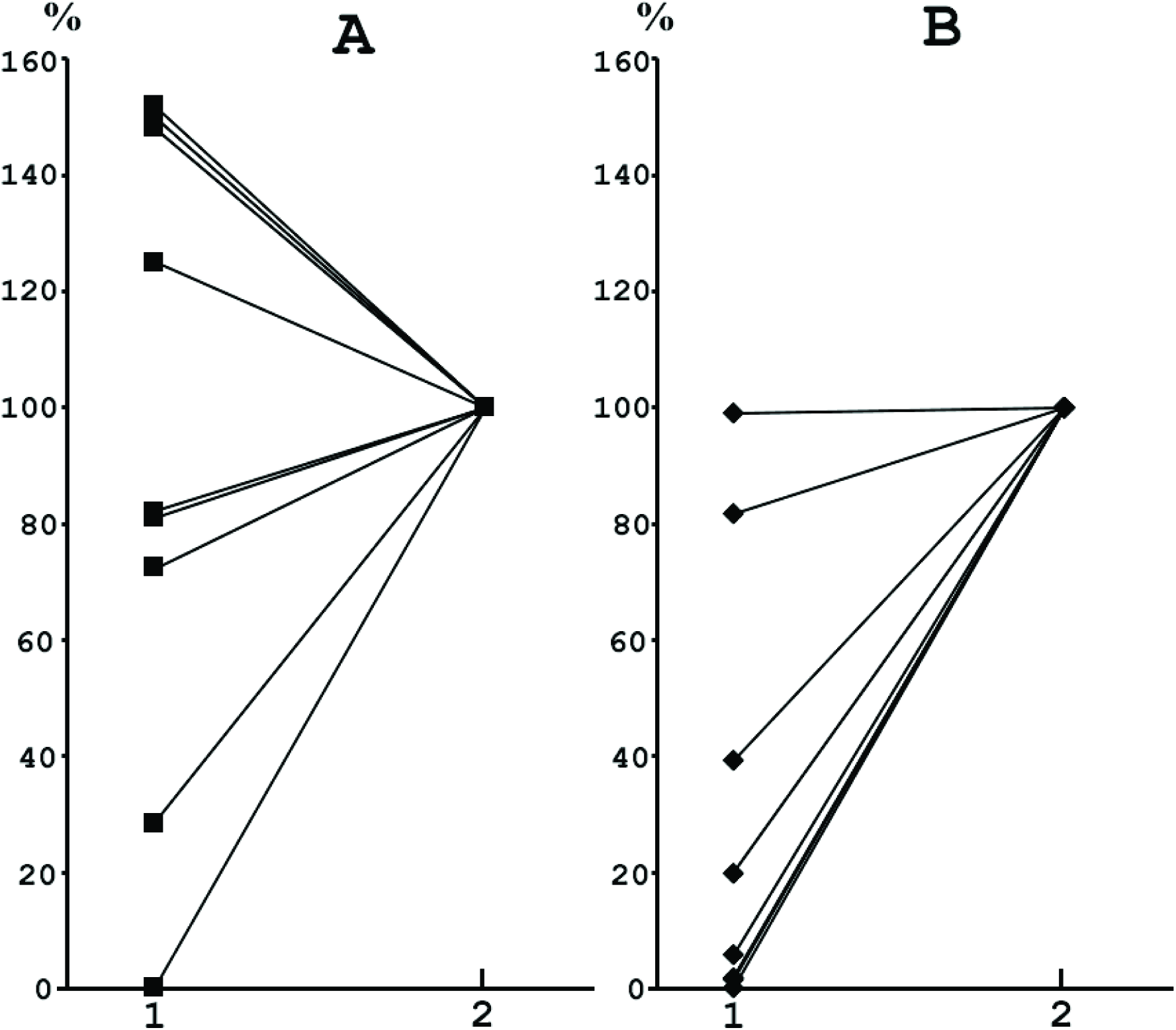
Changes in spontaneous activity level and level of the response to iontophoretic acetylcholine application during hypothermia. The plots schematically represent changes in activity of 9 sensorimotor cortical neurons tested by iontophoretic acetylcholine application during cooling the incubation medium. Electrophoretic acetylcholine current equals 70 nA everywhere (positive pole inside the electrode) A – Average level of spontaneous activity for each neuron. Horizontal axis - temperature values: 1) *t=*26-27°C; 2) *t=*30-31°C; vertical axis – spontaneous firing rate of each neuron as percentage of 30-31°C level taken as 100%. B – Level of response of each neuron to iontophoretic acetylcholine application as a difference between maximal average spike frequency in response periods and in preceding background activity. Horizontal axis – temperature values like on A; vertical axis - level of response of each neuron to acetylcholine as percentage of level of response at *t=*30-31°C taken as 100%.

The presence of two mechanisms regulating the level of spontaneous activity can be revealed by comparison of spontaneous change of activity level under standard temperature, 32-34°C, and under the same temperature before and after hypothermic action. The shift to higher activity level at standard temperature conditions did not lead to significant changes of spike amplitude in cortical neurons (Wilcoxon matched pairs signed rank test, α>>5%). At the same time, the arising hyperactivity in nerve cells registered at the same temperature (32-34°C) after short episodes of hypothermia was almost always accompanied by decrease of spike amplitude (Wilcoxon matched pairs signed rank test, α.< 1%).

For comparison, Fig. 5 shows two cases of increase of spike activity in different neurons during spontaneous change of activity under standard temperature conditions and during post-hypothermic hyperactivity compared to the firing level before cooling. In the first case the increase of firing was not accompanying by changers of spike amplitude (Fig. 5, *A, 1,2*), but in the second case (Fig.5, *B,1,2*) provided by hypothermia raising spontaneous activity 5 minutes after restoration of initial temperature was obtained simultaneously with significant fall of spike amplitude (Wilcoxon – Mann – Whitney test, α<0.005). 10 minutes later in restoration period the level of spontaneous activity decreased to initial value while spike amplitude was still below its meaning before cooling (Fig.5, *B,3*).

**Fig. 5.**
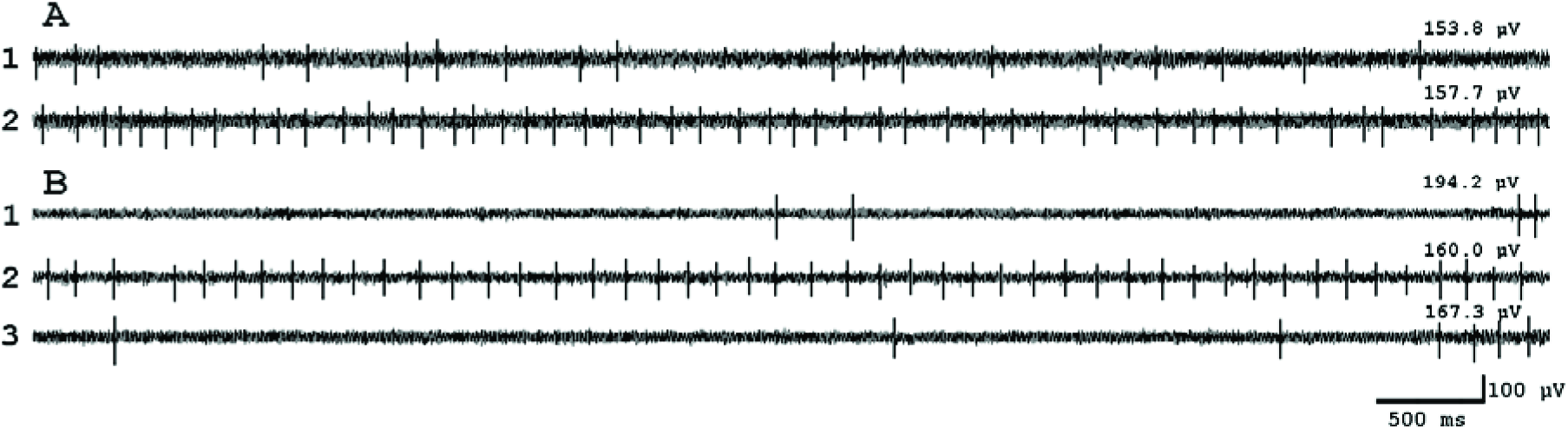
Spontaneous and hypothermia-induced increase of neuronal firing rate. The activity of two different neurons (A and B) is presented. A - Spontaneous increase of spike frequency: 1) and 2) – records of spontaneous firing activity made with 10 min interval. Incubation solution temperature in both cases is 33.2°C. B – Increase of spike activity as a result of hypothermia (cooling to 26°C and further restoration of initial temperature): 1) Activity of the neuron before cooling ( *t*=33.9°C), 2) Activity of the neuron after 10 min hypothermia in 5 min after restoration of initial temperature (*t*=33.4°*C*), 3) Activity of the neuron 15 min after temperature restoration (*t*=33.6°C). Numbers above each records of neuronal activity correspond to average spike amplitude in each fragment.

Thus, the increase of activity frequency actually depends on two mechanisms, one of them being related to the drop of spike amplitude and another being not related to this.

### 3.3. Excitation conducting in dendrites: a regulated parameter during change of spontaneous activity level in standard temperature conditions and after hypothermia.

Both opportunities for increase of spontaneous activity level are realized by more effective action of excitation originating in dendrites onto soma. Glutamate application to particular loci on dendrites evoked the spike reaction registered in soma essentially differ from the reaction to glutamate application directly to soma (Mednikova, Karnup, 1995). The most reliable indicator for these differences is a more prolonged latency period of the spike response during depolarization occurring at dendritic point (Mednikova, Karnup, 1995). The same parameter was the most variable during changes of spontaneous activity level whatever the cause of these changes.

Fig. 6 shows the example of two cases of increasing duration of latency period of response to glutamate applied to dendritic point (100 μm from soma) of the same neuron, when spontaneous activity decreases to zero level. In the first case (Fig.6, *A*), the decrease of frequency took place under influence of cooling the incubation solution to 26°C and lower (Fig.6,*A,4,5*) and it was accompanied by a slight decrease of spike amplitude. In the second case, the frequency dropped spontaneously while registering the activity at constant temperature (near 34°C) 1 hour after hypothermic action (Fig.6,*B*3,4) and almost full restoration of spike activity changed during hypothermia (Fig.6,*B, 1,2*). In both cases, the decrease of spontaneous activity frequency occurred simultaneously with increase of latency period of the response to local glutamate application to dendritic point: from 150 to 400 ms in the first case and from 140 ms to 350 and 700 ms in the second one. Thus, both hypothermia-induced and spontaneous drops of frequency are related to weakening of dendritic impacts onto the neuronal soma.

**Fig. 6.**
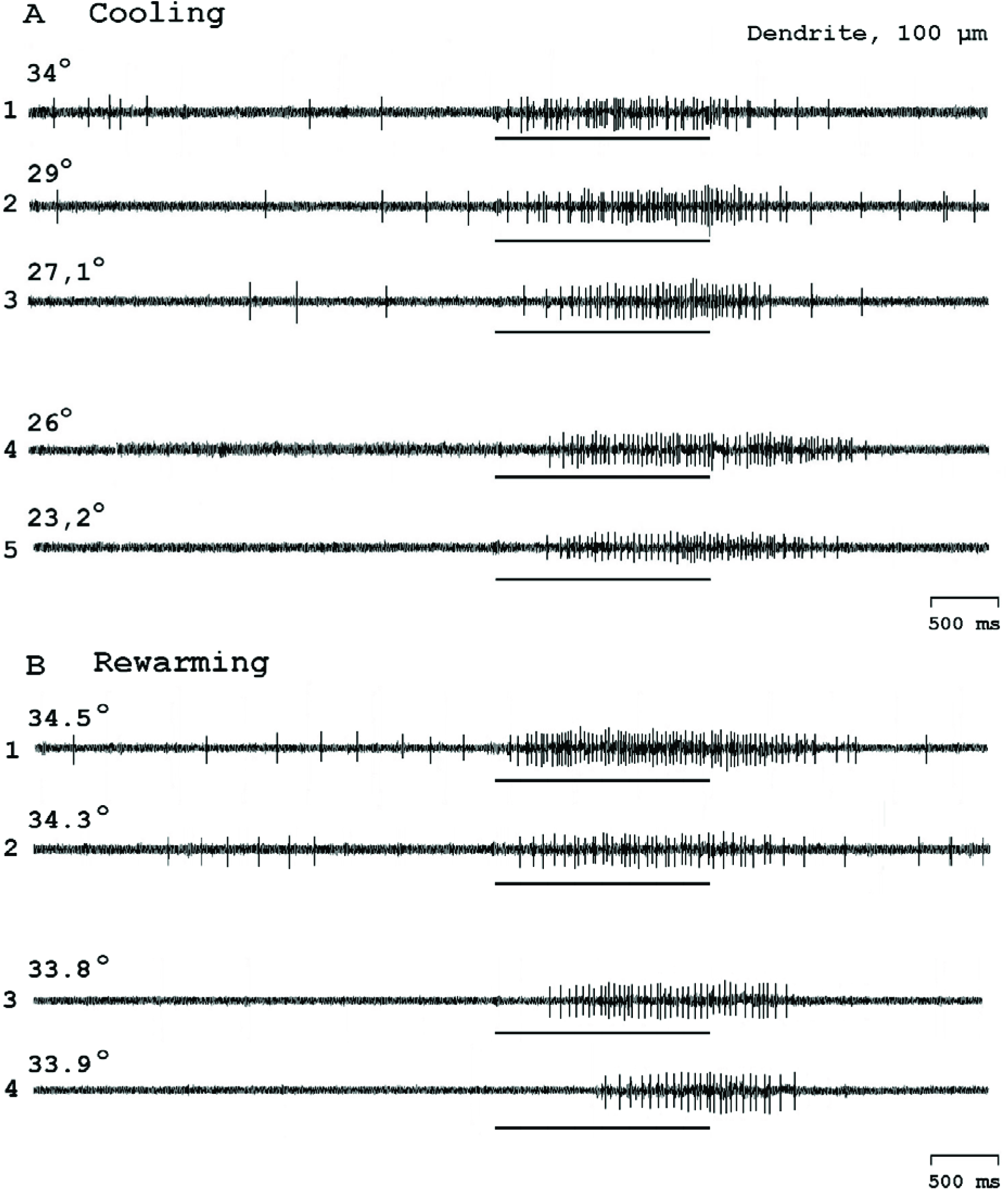
Hypothermia-induced and spontaneous decrease of spike activity of the sensorimotor cortical neuron during simultaneous recording of spike response to iontophoretic glutamate application to dendritic locus. Glutamate was applied to dendritic locus at 100 μm from soma by 80 nA current (negative pole inside the electrode). **A** – activity of the neuron during cooling the incubation medium from 34 to 23.2°C (1-5). **B** – activity of the neuron in post-hypothermic period at different time moments after restoration of initial temperature: 1) 10 min; 2) 15 min; 3) 1 hour; 4) 1 hour 15 min. Time of glutamate iontophoretic current is shown as a line under each record.

Fig. 7 illustrates similar changes under hypothermia happening after the decrease of temperature to 29 °C and below. Spontaneous activity disappears at 29°C (Fig.7, *4*) and increase of latency period of the response to glutamate application to dendritic locus occurs at the same time. The latency period of the reaction reaches the value of 2500 ms at t=28°C (Fig.7, *6*), whereas at t=27°C a reverse process is beginning: the latency period of the response to local dendritic depolarization decreases to 400 ms (Fig.7, *7*), followed by restoration of spontaneous activity with higher frequency at t=28.5°C. Restoration of spontaneous activity is accompanied by decrease of spike amplitude, increase of spike burst activity and masking the activation reaction to glutamate application by intensified spontaneous activity (Fig.7; *8, 9*).

**Fig. 7.**
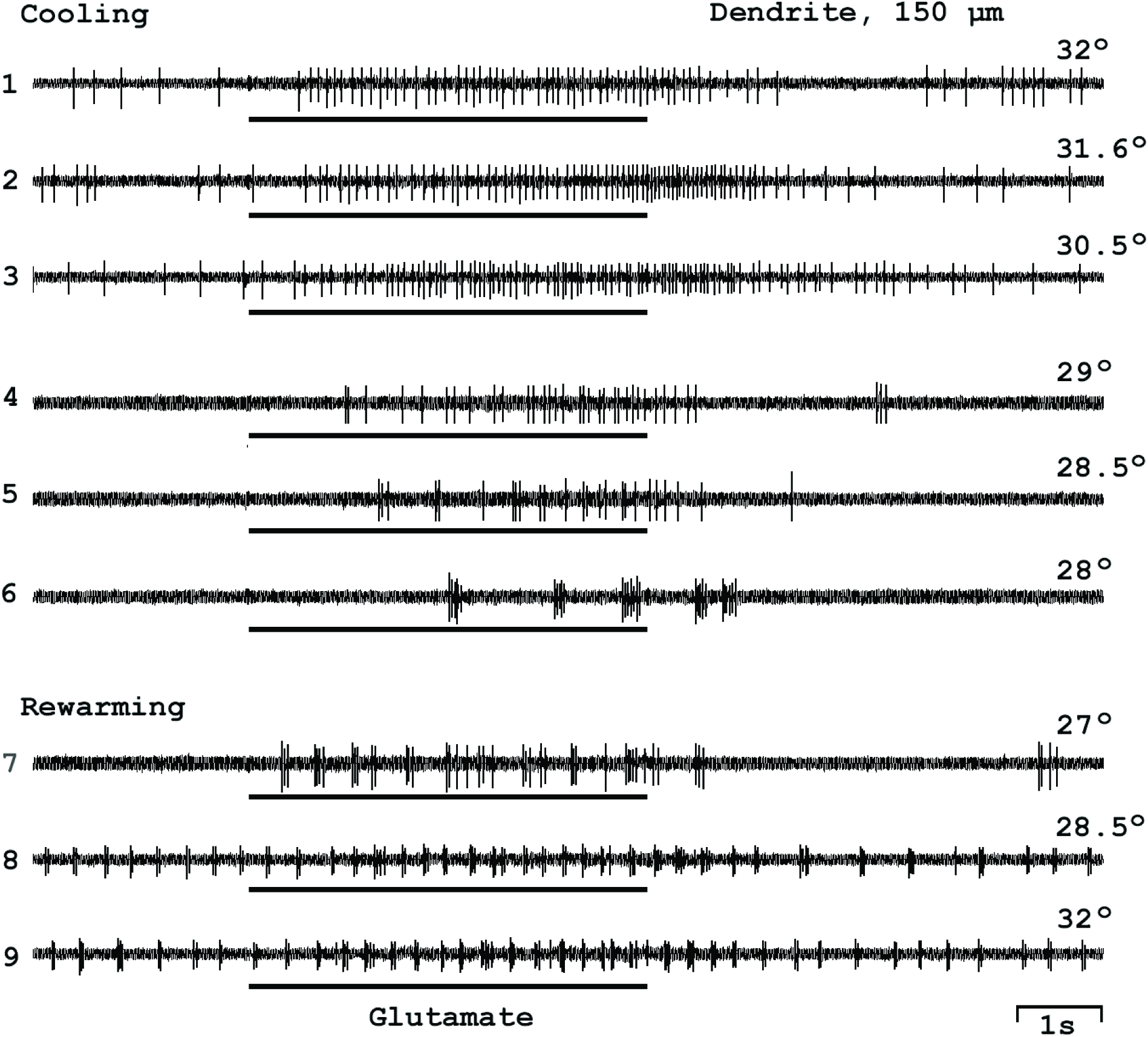
Responses of sensorimotor cortical neuron to iontophoretic glutamate application to dendritic locus during hypothermic decrease of spike frequency and its posthypothermic increase. Glutamate was applied to dendritic locus at 150 μm from soma when cooling the incubation medium from 32 to 28°C (1-6) and during temperature restoration (7-9). Iontophoretic current and duration of its action are the same as for Fig. 6.

Fig. 8 shows a case of appearance of spontaneous activity under hypothermia after decrease of temperature below 26.9 °C in a neuron, which was inactive at higher temperatures. The lifting of spontaneous activity is accompanied by decrease of spike response latency to local dendritic application of glutamate from 650 ms at t=32.8-31.4°C to 350 ms at t=23°C. These changes occur along with decrease of spike amplitude. The following temperature restoration up to initial values demonstrates the hyperactivity phenomenon, i.e. maintenance of spontaneous activity in 10 minutes of incubation above 33°C with simultaneous short-latency reaction to local depolarization of a dendritic point (Fig.8,6). After 15 minutes of restoration period the spontaneous activity disappears and the spike response to dendritic glutamate application simultaneously with spike amplitude are restored (Fig.8, 7).

**Fig. 8.**
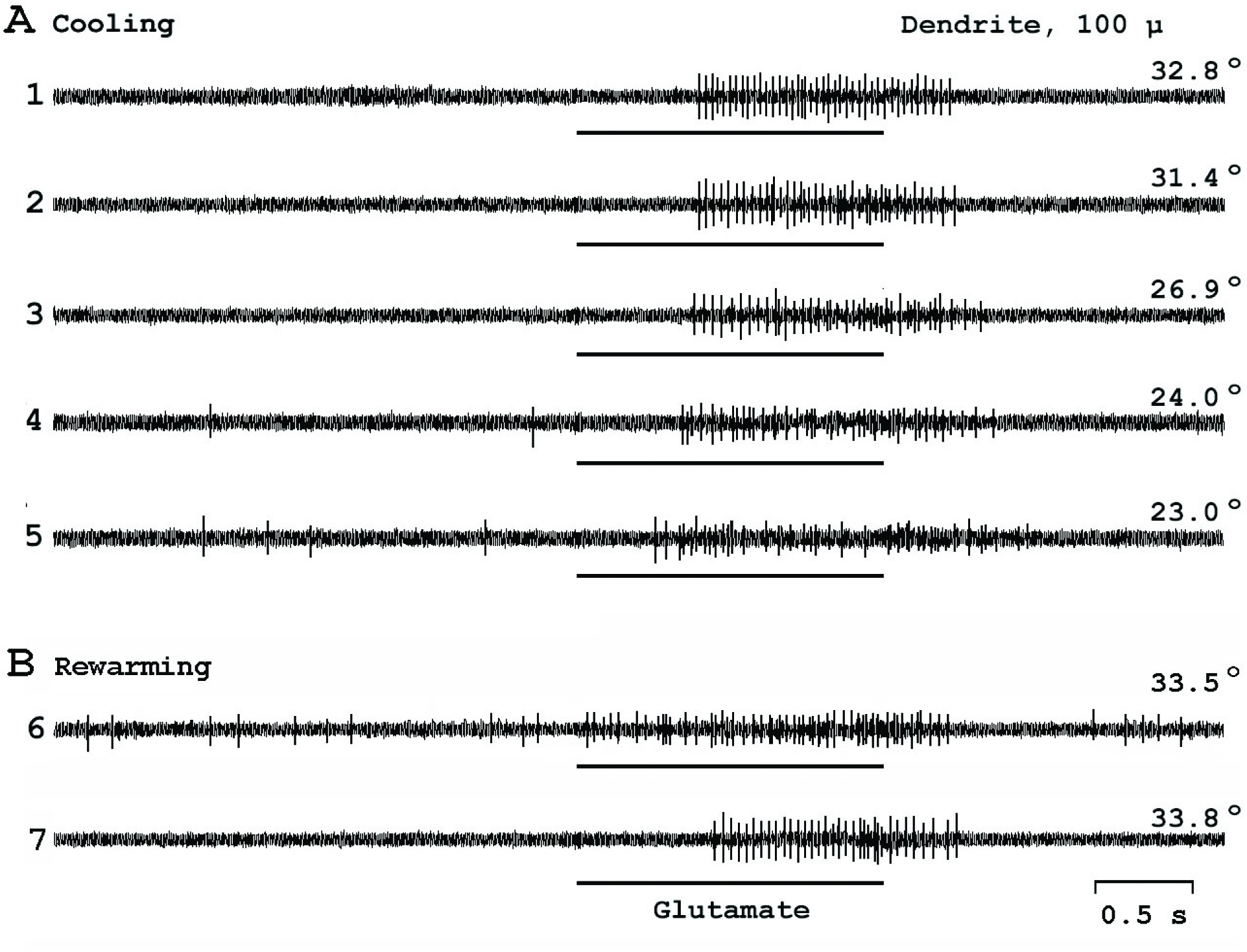
Increase of spontaneous activity of the neuron during hypothermia and responses to glutamate application to dendritic locus._ Glutamate was applied to dendritic locus at 100 μm from soma. **A**) Records of neuronal activity during cooling the incubation medium from 32.8 to 23°C(1-5) **B**) Records of neuronal activity during restoration of incubation solutiontemperature: 6-5 min after restoration, 7- 15 min after restoration. Iontophoreticcurrent and duration of its action are the same as for Fig. 6.

Stimulation of dendritic points (distance from soma ranging from 50 to 250 μm) under hypothermic conditions was carried out on 20 neurons. As well as spontaneous activity level, the responses induced from dendrites also changed during cooling. Decrease of spontaneous activity level (6 neurons) was always accompanied by increase of latency periods or even disappearing of the response, independent on distance from the soma. Increase of spike frequency when cooling or during post-hypothermic period took place either along with decrease of latency period of the response to dendritic stimulation (6 neurons), or (in the cases of markedly increase of spike frequency) along with masking of the response (3 neurons). If hypothermia did not cause changes in spontaneous activity, the changes in response expression to glutamate applied to dendrites also lacked (5 neurons).

On the contrary spike responses to glutamate applied to soma (6 neurons) were always very stable independently of the level of spontaneous activity. Latency periods were constant, ranging from 30 to 400 ms in different neurons. In cases of increase or decrease of spike frequency under hypothermic conditions, the response to glutamate proportionally increased (4 neurons) or decreased (1 neuron). When the spike frequency increased significantly, masking of the response was detected (1 neuron).

Thus, despite all the diversity of symptoms, the analysis of the effects of hypothermia on the activity of cortical neurons revealed the following:

1) A correlation between alteration of the level of spontaneous activity caused by hypothermia and the efficiency of conductive function of dendrites was found.

2) In most of the cases, the decrease of spike amplitude occurred when temperature was decreased.

3) The revealed changes in the activity of cortical nerve cells occurred in temperature interval from 27 to 29°C, where excitatory spike reactions to acetylcholine sharply attenuates.

## 4. Discussion

The data obtained give evidence that hypothermic action leads to serious disorders in the activity of cortical nerve cells. Small changes are occurring even in 30-34°C temperature range, comprising slight decrease of spike amplitudes of cortical neurons, but global problems begin to take place when cooling below 30°C. As shown earlier (Mednikova et al., 2008) and confirmed by the current study (Fig.3 и Fig.4), below this point, there is the first transition zone for M-cholinergic brain reaction, where (27-29°C) the decrease of its rate occurs. M-cholinergic process blocks K^+^ channels on neuronal membranes (Brown et al.,1995; Krnjević et al.,1971; McCormick D.A., Prince D.A., 1986) and its sharp temperature related decrease provides additional growth of K^+^ permeability and facilitates K^+^ efflux from nerve cells in the direction of concentration gradient. Such ionic flow is so significant in mammals that the capabilities of Na^+^, K^+^-ATPase to restore ionic homeostasis cannot fully compensate the disorder. The membrane potential begins to decrease, which leads to significant decrease of spike amplitude even under short hypothermia action (Fig.2). All other hypothermia-induced disorders are the consequences of these two processes, i.e. temperature limitation of M-cholinergic reaction rate and progressive K^+^ ion accumulation in extracellular space.

The most varying parameter during cooling is the frequency of spontaneous spike activity in nerve cells. The basic reason for formation of spontaneous activity is the constant flow of miniature dendritic glutamatergic EPSPs (Williams, Stuart, 2002), which can have an amplitude of several tens mV at the origin points on dendrite loci (MacGregor, 1968), but are rapidly attenuated during motion to the soma (Williams, Stuart, 2002). The conditions affecting more effective excitation conduction along dendrites are determined by geometry of the neurons and specific resistance of their membranes (Rall et al.,1992). This is why acetylcholine, blocking K^+^ permeability of the neuronal membranes when interacting with M-cholinoreceptors (Brown et al.,1995; Krnjević et al.,1971; McCormick D.A., Prince D.A., 1986), leads to both more effective excitation conduction along the dendrites (Mednikova et al. 1998) and to increase of spontaneous activity level (Mednikova et al., 2010).

Limitation of K^+^ permeability of the neuronal membranes could also be achieved via increase of extracellular K^+^ concentration, which takes place at temperature below 27°C, when the rate of M-cholinergic process decreases. As a result, the specific resistance of cell membranes is increasing, which has been recorded as increase of input resistance below 27°C (Aihara et al. 2001; Thompson et al., 1985) and has as consequence more effective conducting the flow of miniature excitatory postsynaptic potentials from dendrites to soma for its transformation into spike process. The proof for large increase of extracellular concentration of K^+^ ions is the increase of spike width and slowing of rates of spike rise and fall below 27 °C recorded intracellularly (Aihara et al. 2001; Thompson et al., 1985), as well as bursts formation in activity of some neurons after cooling and further temperature restoration (Fig.7). The data adduced point on decrease of repolarization rate of neuronal membranes after depolarization potential shifts, which is the consequence of increased K^+^ concentration in extracellular medium. Therefore we conclude that two processes occur under hypothermia: from one side, the increase of membrane K^+^ permeability via decrease of M-cholinergic reaction rate at 27-29°C, and from another side, decrease of K^+^ permeability due to progressive growth of K^+^ concentration in extracellular space. Prevailing of one or another process should depend on structural and membrane properties of the neurons, which would determine the dynamics of spontaneous spike activity changes of cortical neurons under cooling (Fig.1).

Density of K^+^ channels on the membranes of mammalian cortical neurons additionally increases after the birth of animals (Kang et al.,1996), exceeding 3-5 times to 28^th^ day of postnatal development the level at the first day of life. The increase of K^+^ channel density is accompanied by increase of the diversity of their expression on the membranes of various cortical neurons (Kang et al., 1996). Thus, the density of K^+^ channels in mammalian neuronal membranes is different from cell to cell. Due to this, one can imply that neurons having low initial spontaneous activity level (up to 4 impulses per second, i. e. groups I and II) have a large average density of K^+^ channels in the membranes. It may determine their low activity level as a result of a large decrement of EPSP amplitude when moving along the dendrites. For the same reason, hypothermic increasing K^+^ permeability of the membranes below 27°C leads to rapid increase of K^+^ concentration outside of these cells. That is why fast decrease of spike amplitude occurs, whereas spontaneous activity was increasing with further accumulation of K^+^ ions in extracellular environment as a result of concentration-dependent increase of membrane resistance (Fig.1, *A,I,II*).

The opposite effect caused by hypothermia mainly in neurons with activity above 4 spikes per second (groups III, IV) is apparently linked to relatively low content of K^+^ channels in their membranes. As a result, under 27 °C the prevailing influence on spontaneous activity level is connected with opening of K^+^ channels, leading to drop of conductive function of the dendrites and decrease of activity rate. Increase of extracellular K^+^ concentration under these conditions is slow, appearing as a small decrease of spike amplitude and a slight hyperactivity after temperature restoration.

Efflux of K^+^ ions from neurons beginning at 27 °C does not end further and can lead to a significant decrease of membrane potential level under further cooling or just in a certain time course (Aihara et al. 2001). Against depolarization shift, the drop of spike amplitude takes place (Fig.2), which is confirmed by earlier investigations (Mednikova et al.,2002; Mokrushin et al., 2014; Volgushev et al.,2004) and the destructive intracellular processes are triggered (Hochachka, 1986; Ivanov, 2004). But even a small depolarization for potential-dependent K^+^ channels is quite enough to open ever more (Adams et al. 1982), causing further increase of K^+^ efflux from the cell and further depolarization shift (Aslanidi, 1997). Thus a self-maintained process appears, resulting in increased extracellular K^+^ concentration maintained for a long time after restoration of initial brain temperature. Experimentally it was observed as a long period of increased posthypothermic spontaneous activity frequency and long period of the decrease of spike amplitude.

The lack of change in activation spike reactions during glutamate application to soma on cooling proves the results of previous studies (Mednikova et al.,2002) and evidences that glutamate sensitivity is not changing under hypothermia. This fact allows us to draw a conclusion that whatever the reason of change of neuronal spontaneous activity would be, its rate is finally related to the decrement of glutamatergic excitation moving along the dendrites. Fig. 6 and 7 show that the disappearing of spontaneous activity below 27-29°C is coupled to prolongation of latency periods of responses to iontophoretic glutamate application to dendritic loci (Fig.6,*A,4,5* and Fig. 7,*4,5,6*). This means that under hypothermia the excitation passing along the dendrites is so strongly attenuated that it can hardly penetrate the soma, overcoming its shunting properties. The same event can happen spontaneously (Fig.6, *B, 3, 4*), without altering temperature, but due to spontaneous limitation of acetylcholine efflux from cholinergic synapses in the cortex (Houser et al.,1985) in registered zone. During the decrease of spontaneous activity under stationary conditions the prolongation of latency period of the response to glutamatergic stimulation of the dendrite appears even more significant than under action of hypothermia (Fig.6,B,4 and 6,A,5), because it is regulated only by attenuation of influence from cholinergic system and is not interfered by the increase of membrane resistance related to the increase of extracellular K^+^ content. Thus, the dendrites should be considered as special machinery for spontaneous activity formation, which is finely regulated via alteration of K^+^ permeability of neuronal membranes. At the temperatures specific for homoeothermic animals in stationary conditions it occurs mainly due to the regulating effect of brain cholinergic system.

The rate of M-cholinergic reaction in temperature range 27-36°C is sharply increasing twice. The first temperature-dependent shift at 27-29°C is a reason for hypothermic disorder of activity of the cortical neurons. The second temperature-dependent shift occurs at 34-36°C and is characterized by significant increase of the rate of the process, its increment being on average for different neurons 200% per 1 degree compared to the 12-13% per degree in the fist temperature zone (Mednikova et al., 2008). The high rate of cholinergic reaction regulating the permeability state of K^+^-channels of neuronal membranes at brain temperatures normal for homoeothermic animals requires great energetic support. The energetic demands of this process can be full provided only by perfecting of circulatory, respiratory, digestive and other organs, which makes the difference between the any of homoeothermic animal and their poikilothermic evolutionary precursors (Ivanov, 2004). Thermodynamically almost impossible affords to regulate specific for homoeothermic animals low-ohm neuronal membranes lead to high dependence of their brain from energetic supply, the lack of which leads straight to hypoxic disorders (Hochachka 1986; Ivanov 2004; Lipton, 1999). The alterations caused by hypoxia are absolutely identical to processes occurring under hypothermia because the first sign of hypoxia is limitation of Na^+^, K^+^-ATPase activity followed by accumulation of K^+^ ions in extracellular space (Lipton, 1999). That is why hypoxia and hypothermia are equally destructive for mammals, and Nature leaves us only 1.5 °C temperature interval for normal functional work of our perfect brain (Ivanov, 2004).

## 5. Conclusion

The main reason of hypothermic disorders in functional activity of cortical neurons is temperature-dependent limitation of the rate of M-cholinergic reaction of the brain, which is a regulator of membrane properties of nerve cells by closing K^+^ channels. The decrease of this process rate below 27°C reduces the influence of dendritic excitation on the soma and in the case of homoeothermic animals leads to neuron homeostasis disorders and K^+^ accumulation in extracellular space. This is the reason of spike amplitude attenuation and pathological rise of spontaneous firing of cortical neurons in hypothermic conditions. Different functional properties of cortical neurons determine the peculiarities of their response to hypothermia.

## Acknowledgements

The work was supported by the Russian Foundation for Basic Research (project number 13-04-01114a)

